# Primary Target Prediction of Bioactive Molecules from Chemical Structure

**DOI:** 10.1101/413237

**Authors:** Abed Forouzesh, Sadegh Samadi Foroushani, Fatemeh Forouzesh, Eskandar Zand

## Abstract

There are various tools for computational target prediction of bioactive molecules from a chemical structure in a machine-readable material but these tools can’t distinguish a primary target from other targets. Also, due to the complex nature of bioactive molecules, there has not been a method to predict a target and or a primary target from a chemical structure in a non-digital material (for example printed or hand-written documents) yet. In this study, an attempt to simplify primary target prediction from a chemical structure was resulted in developing an innovative method based on the minimum structure which can be used in both formats of non-digital and machine-readable materials. A minimum structure does not represent a real molecule or a real association of functional groups, but is a part of a molecular structure which is necessary to ensure the primary target prediction of bioactive molecules. Structurally related bioactive molecules with the minimum structure were considered as neighbor molecules of the query molecule. The known primary target of the neighbor molecule is used as a reference for predicting the primary target of the neighbor molecule with an unknown primary target. In results, we confirmed the usefulness of our proposed method for primary target prediction in 548 drugs and pesticides involved in four primary targets by eight minimum structures.

## INTRODUCTION

Bioactive molecules such as drugs and pesticides are produced in large numbers by many commercial and academic groups around the world^18^. Most bioactive molecules perform their actions by interacting with proteins or other macromolecules^5^. However, for a significant fraction of them, the primary target remains unknown^5^.

Computer-aided drug design, that offers an *in silico* alternative to medicinal and agricultural chemistry techniques for studying the structure and predicting the biological activity of drug and pesticide candidates, has the advantages of both speed and low cost and is becoming an indispensable program of major pharmaceutical and agrochemical companies^6^.

With the ever-increasing public availability of bioactivity data^2^, it is possible to construct reliable target prediction models using statistical or machine learning methods^21^. However, bioactivity data hasn’t been increased in all areas. For example, the databases are rich in human targets and molecules that modulate these targets, but contain limited information when it comes to bacterial targets^11^.

Various methods (such as chemical structure similarity searching^9^, data mining/machine learning^17^, panel docking^14^ and bioactivity spectra based algorithms^3^) can be employed for computational target prediction (each method adopts a particular model type). The tools created by these methods have not been able to distinguish between a primary target and other targets yet due to the focus of databases from which they have been extracted. Also, they are not equally reliable in all areas. Furthermore, chemical structures should be provided in machine-readable descriptions so that the tools can be used.

Based on our knowledge, there is no method to predict a target and or a primary target from a chemical structure in a non-digital material. In this study, an attempt to simplify primary target prediction from a chemical structure was resulted in developing an innovative method based on the minimum structure which can be used in both formats of non-digital and machine-readable materials. The proposed method has several distinctive features compared to available computational target prediction methods. Firstly, it’s easy to use. Secondly, it is highly accurate. Thirdly, it can be used appropriately in both formats of non-digital and machine-readable materials. Fourthly, it enables us to gain a deeper understanding (more informative) of the relationship between the chemical structure and the primary target.

## METHODS

The proposed method steps for primary target prediction of bioactive molecules from a chemical structure include (i) query molecule, (ii) similarity searching, (iii) data collection, (iv) minimum structure identification and (v) primary target prediction (example in Supplementary Figure 9).

### (I) Query molecule

A bioactive molecule such as a drug or a pesticide is used as a query molecule. The query molecule may have a known or an unknown primary target. The query molecule with known primary target can be used as a reference to predict the primary target of structurally related bioactive molecules.

### (II) Similarity searching

The query molecule is used to search for the structurally related bioactive molecules with similar chemical scaffold. However, it must be born in mind that structurally related analogs may bind in a slightly or considerably different manner^24^. KEGG^7^, DrugBank^23^, PubChem^10^ and ChEMBL^4^ databases provide common names and chemical structures for large numbers of bioactive molecules and, in some cases, their primary targets. All four databases support structure similarity searches.

### (III) Data collection

The primary target information is collected for all structurally related bioactive molecules. If the primary target of the query molecule is not known, information on structure-activity relationship and pharmacophore will be collected for all structurally related bioactive molecules. If the primary target of the query molecule is known, information on structure-activity relationship and pharmacophore will be collected only for structurally related bioactive molecules with the same primary target as the query molecule.

Information on the primary target, structure-activity relationship and pharmacophore are obtained from databases with annotated primary target (such as KEGG^7^, DrugBank^23^, PubChem^10^ and ChEMBL^4^ databases), scientific literature and pharmacophoric descriptors (including hydrogen bonds, hydrophobic and electrostatic interaction sites). The most recent development regarding pharmacophore alignment technique is LigandScout’s pattern matching approach^25^.

### (IV) Minimum structure identification

A minimum structure does not represent a real molecule or a real association of functional groups, but is a part of a molecular structure which is necessary to ensure the primary target prediction of bioactive molecules. The minimum structure is identified using data collection about the structurally related bioactive molecules. The minimum structure consists of the core and or the peripheral part. Here, the peripheral part is shown as the comment. The core plays an essential role in a bioactive molecule. Furthermore, modifying at some key position on the peripheral part can make a big change in the primary target or the activity of a bioactive molecule. Thus, the peripheral part can be useful for distinguishing the bioactive molecules based on their primary targets.

Since the minimum structure is related to the structurally related bioactive molecules and information about them, when they become available, the minimum structure can be updated to further refine it.

### (V) Primary target prediction

Structurally related bioactive molecules with the minimum structure were considered as neighbor molecules of the query molecule. The known primary target of the neighbor molecule is used as a reference for predicting the primary target of the neighbor molecule with an unknown primary target.

## RESULTS AND DISCUSSION

In results, we made predictions for eight groups of bioactive molecules. Here, the proposed method was employed in 548 drugs and pesticides involved in four primary targets (Tables 1-4 and Supplementary Data 1). 4-Pyridone group includes 192 bioactive molecules of DNA gyrase and topoisomerase IV inhibitors (Table 1; example in Supplementary Figure 1), 2,4(or 5)-diaminocyclohexanol group includes 138 bioactive molecules of small ribosomal subunit inhibitors (Table 2; example in Supplementary Figure 2), (4a*RS*,5a*RS*)-Sancycline group includes 34 bioactive molecules of small ribosomal subunit inhibitors (Table 2; example in Supplementary Figure 3), cytosine group includes 28 bioactive molecules of large ribosomal subunit inhibitors (Table 3; example in Supplementary Figure 4), 3-glutarimidyl group includes 17 bioactive molecules of large ribosomal subunit inhibitors (Table 3; example in Supplementary Figure 5), (1*R*)-propanol group includes 10 bioactive molecules of large ribosomal subunit inhibitors (Table 3; example in Supplementary Figure 6), imidazol-1-yl group includes 54 bioactive molecules of sterol 14α-demethylase inhibitors (Table 4; example in Supplementary Figure 7), and 1,2,4-triazol-1-yl group includes 75 bioactive molecules of sterol 14α-demethylase inhibitors (Table 4; example in Supplementary Figure 8).

**Table 1.**
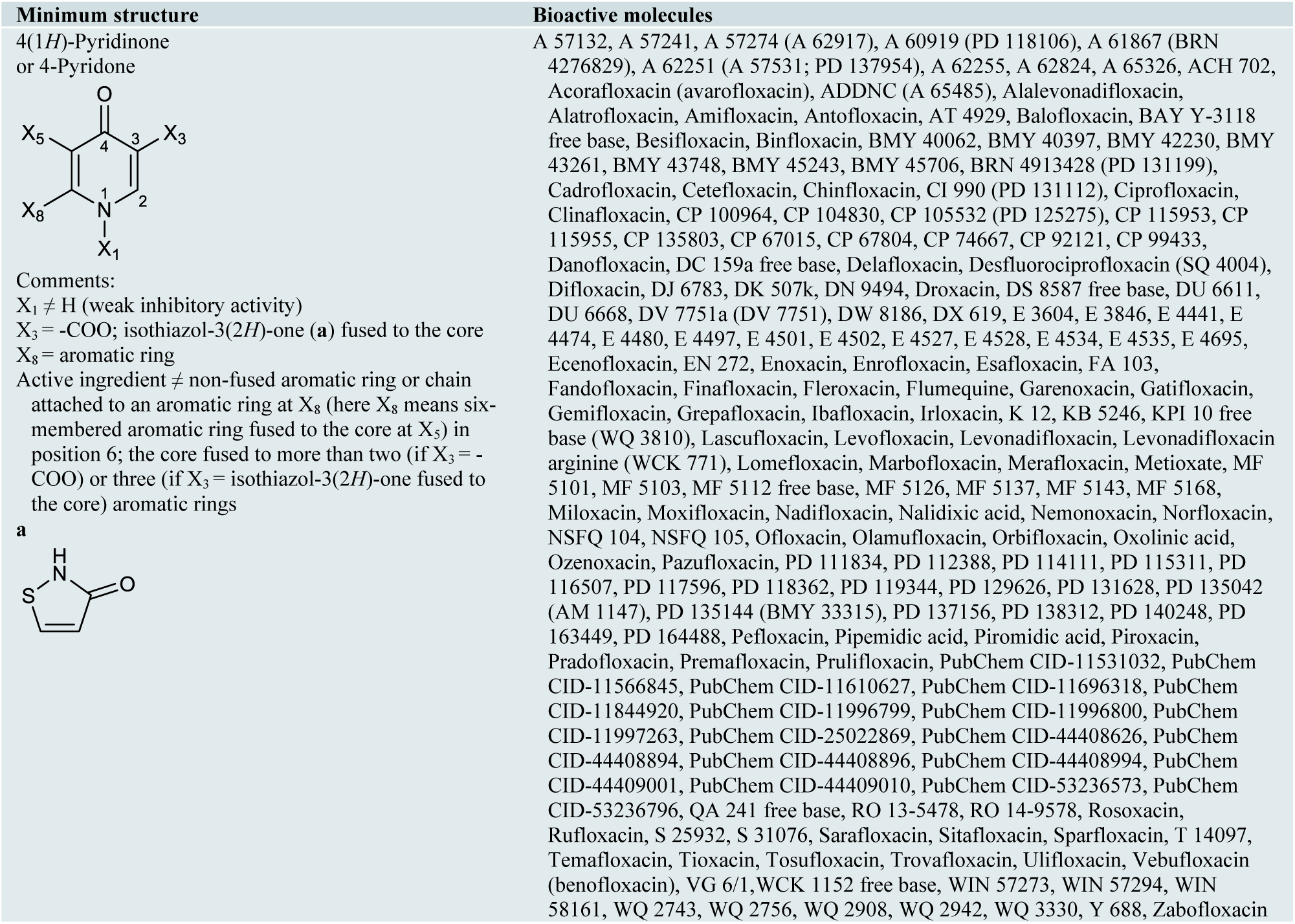
The bioactive molecules with primary target of bacterial type IIA topoisomerase (DNA gyrase and topoisomerase IV) inhibition predicted by the minimum structure.

**Table 2.**
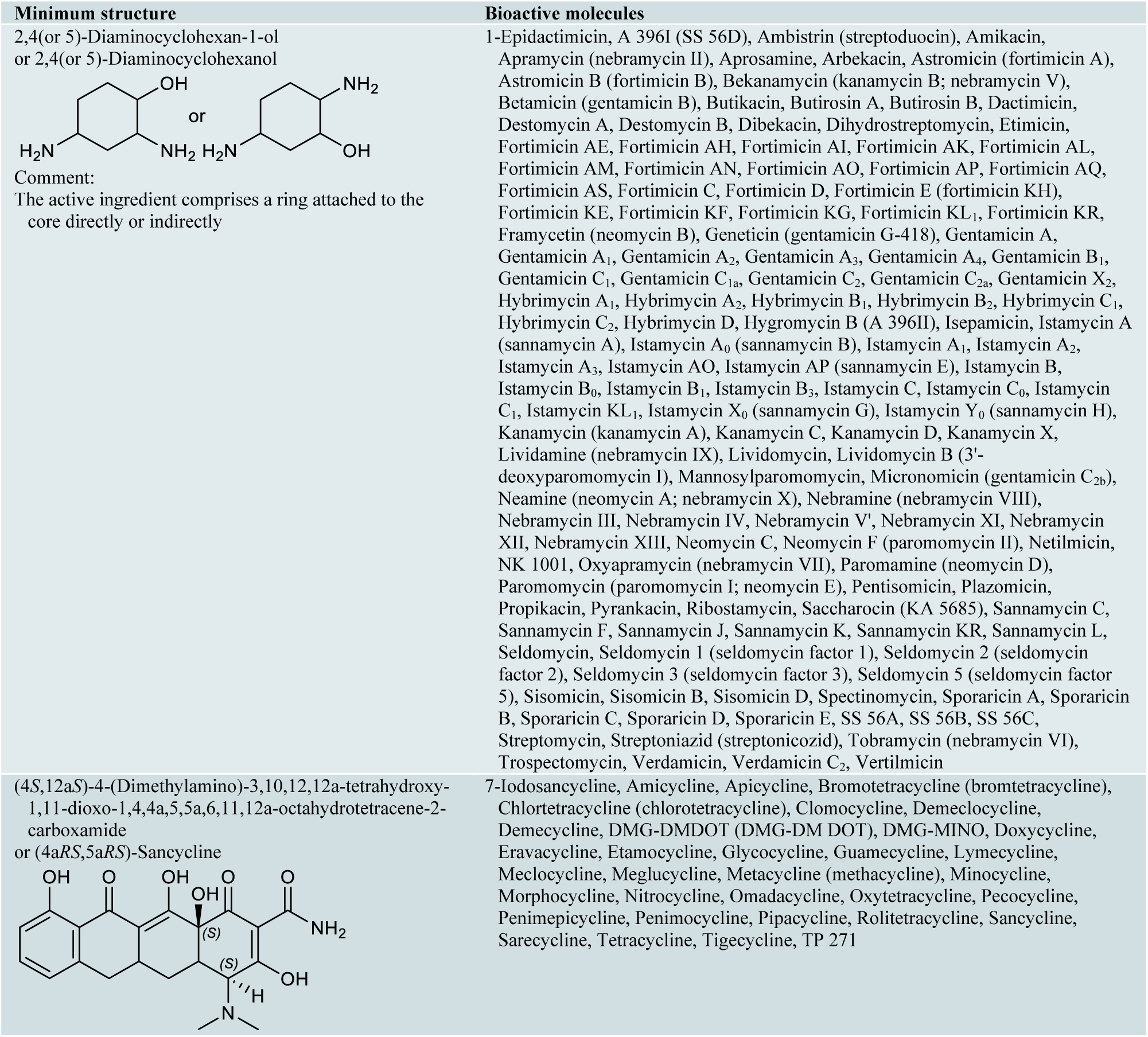
The bioactive molecules with primary target of small ribosomal subunit inhibition predicted by the minimum structure.

**Table 3.**
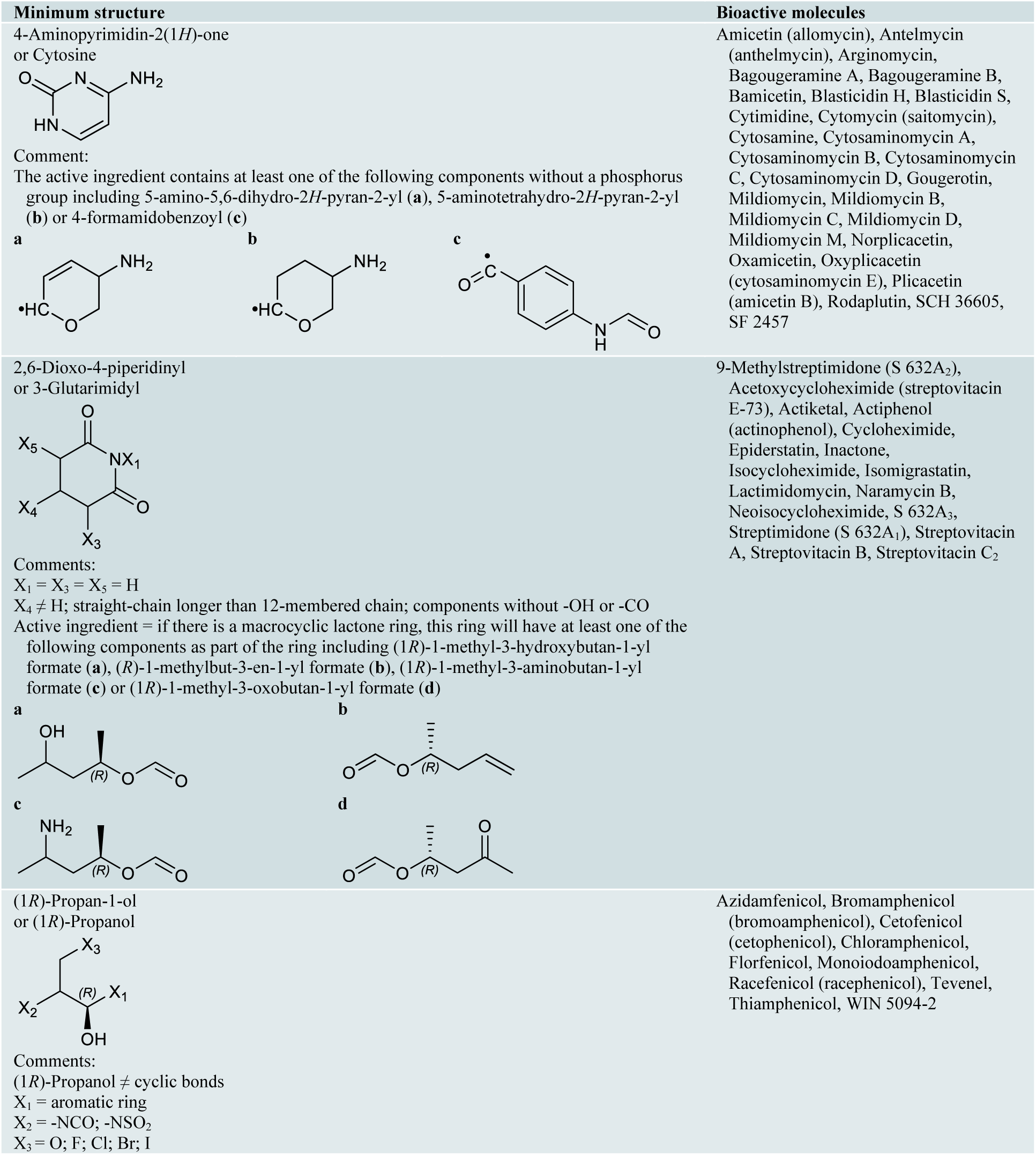
The bioactive molecules with primary target of large ribosomal subunit inhibition predicted by the minimum structure.

**Table 4.**
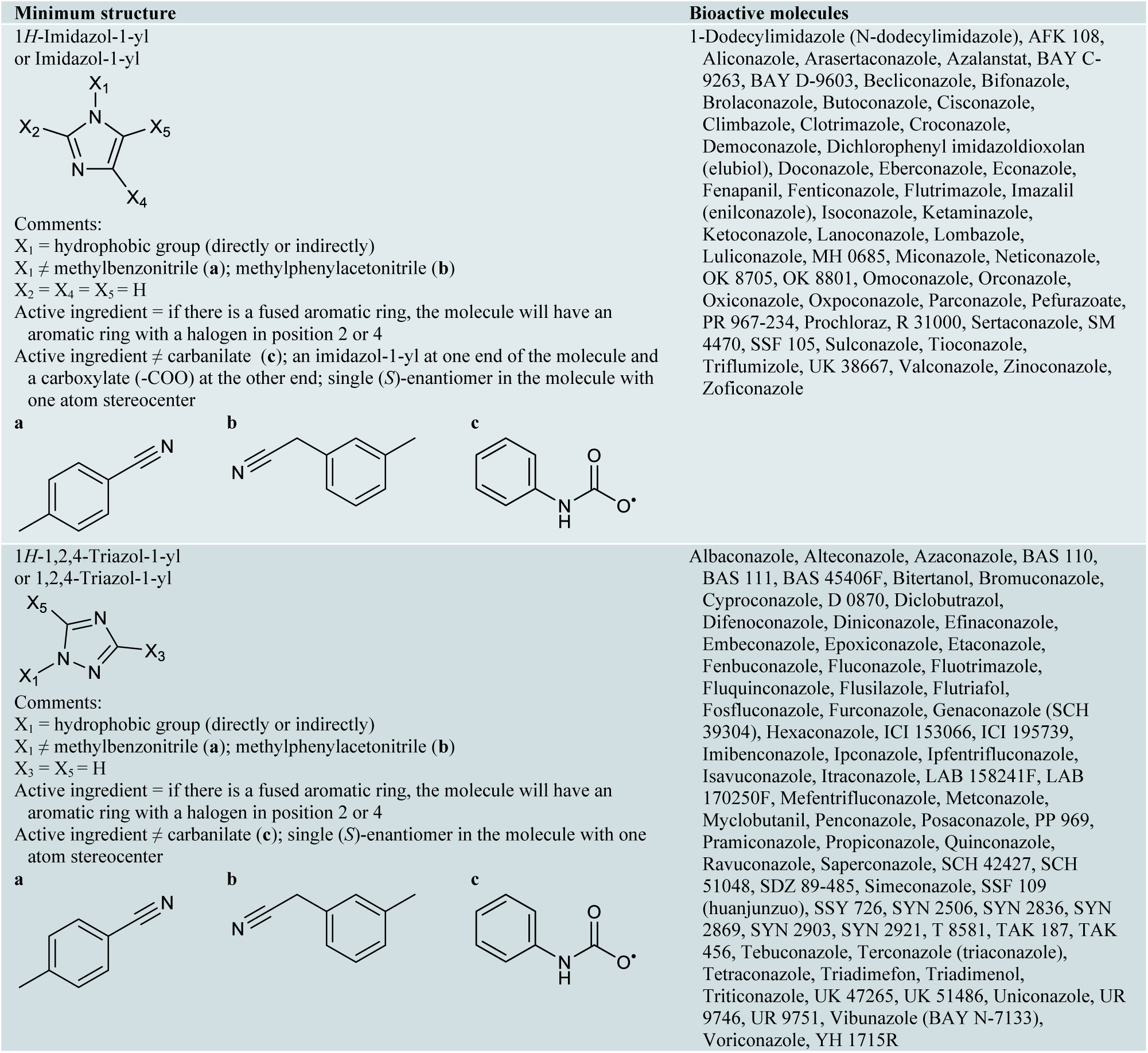
The bioactive molecules with primary target of sterol 14α-demethylase inhibition predicted by the minimum structure.

Many of these 548 predictions were either confirmed by scientific literature, database searching or computational target prediction tools. Out of 548 predictions, 371 predictions (67.7%) were confirmed by scientific literature published in scientific journals, conferences and books (4-pyridone group with 24.5%, 1,2,4-triazol-1-yl group with 11.3%, 2,4(or 5)-diaminocyclohexanol group with 11.3%, imidazol-1-yl group with 6.9%, (4a*RS*,5a*RS*)-Sancycline group with 4.7%, cytosine group with 4.6%, 3-glutarimidyl group with 2.7% and (1*R*)-propanol group with 1.6%; Supplementary Data 2), 160 predictions (29.2%) were confirmed by database searching (KEGG^7^, PubChem^10^, DrugBank^23^ and ChEMBL^4^ respectively with 23.2%, 16.4%, 11.3% and 10.4%; Supplementary Data 3) for annotated primary target, 546 predictions (99.6%) were confirmed by computational target prediction tools (PASS online^13^ with 99.3%, SEA^8^ with 58%, PPB^1^ with 54.4%, TargetHunter^20^ with 44.7%, ChemProt^12^ with 44.5%, PharmMapper^22^ with 37.8%, SuperPred^16^ with 8.6%, HitPick^15^ with 6.4% and SPiDER^19^ with 0.4%; Supplementary Data 4). We found no precedent for one prediction.

A lot of information is needed to identify the minimum structure because it is necessary to know which part(s) of the bioactive molecule confers the activity. Therefore, the proposed method leads to a high accuracy and a deeper understanding of the relationship between the chemical structure and the primary target. Furthermore, after identifying the minimum structure, it’s been easy to use and it can be used in both formats of non-digital and machine-readable materials. Due to the complex nature of bioactive molecules, there has not been a method to predict a target and or a primary target from a chemical structure in a non-digital material yet and the proposed method is the only method that can be used for this purpose.

The proposed method is not applicable in cases where no neighbor bioactive molecules for a primary target exist, since in these situations no training on the minimum structure-based information is possible.

---

**Supplementary Figure 1.**
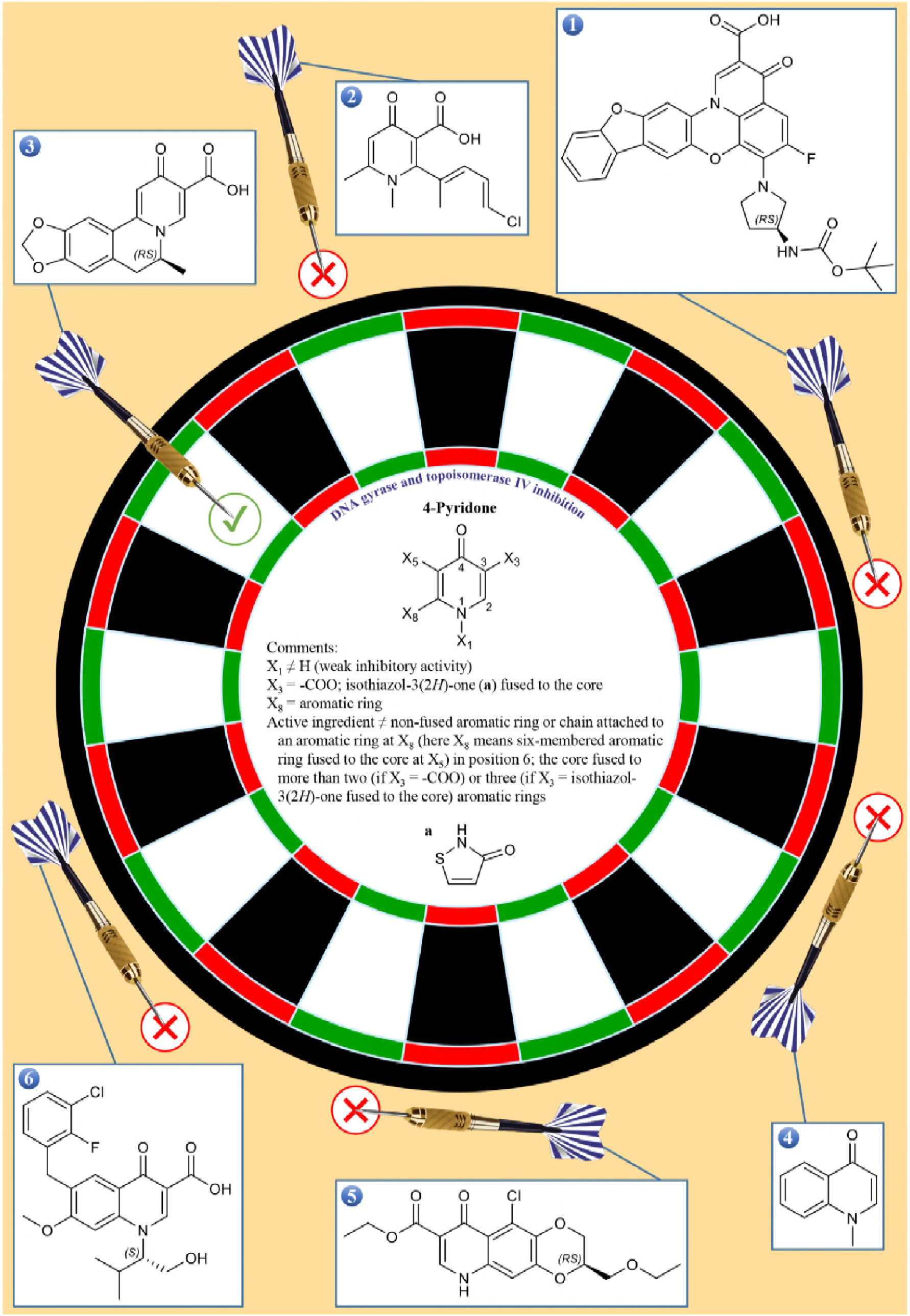
An example of identifying a bioactive molecule with the primary target of DNA gyrase and topoisomerase IV inhibition by the minimum structure in 4-pyridone group. Molecule 1 contains five aromatic rings fused to the core (4-pyridone), molecule 2 contains one methyl at X_8_, molecule 4 doesn’t contain -COO or isothiazol-3(2*H*)-one fused to the core at X_3_, molecule 5 contains hydrogen at X_1_ and molecule 6 contains one methyl attached to the aromatic ring (phenyl) at X_8_ (here X_8_ means six-membered aromatic ring fused to the core at X_5_) in position 6. Therefore, the primary target of molecules 1, 2, 4, 5 and 6 is not DNA gyrase and topoisomerase IV inhibition.

**Supplementary Figure 2.**
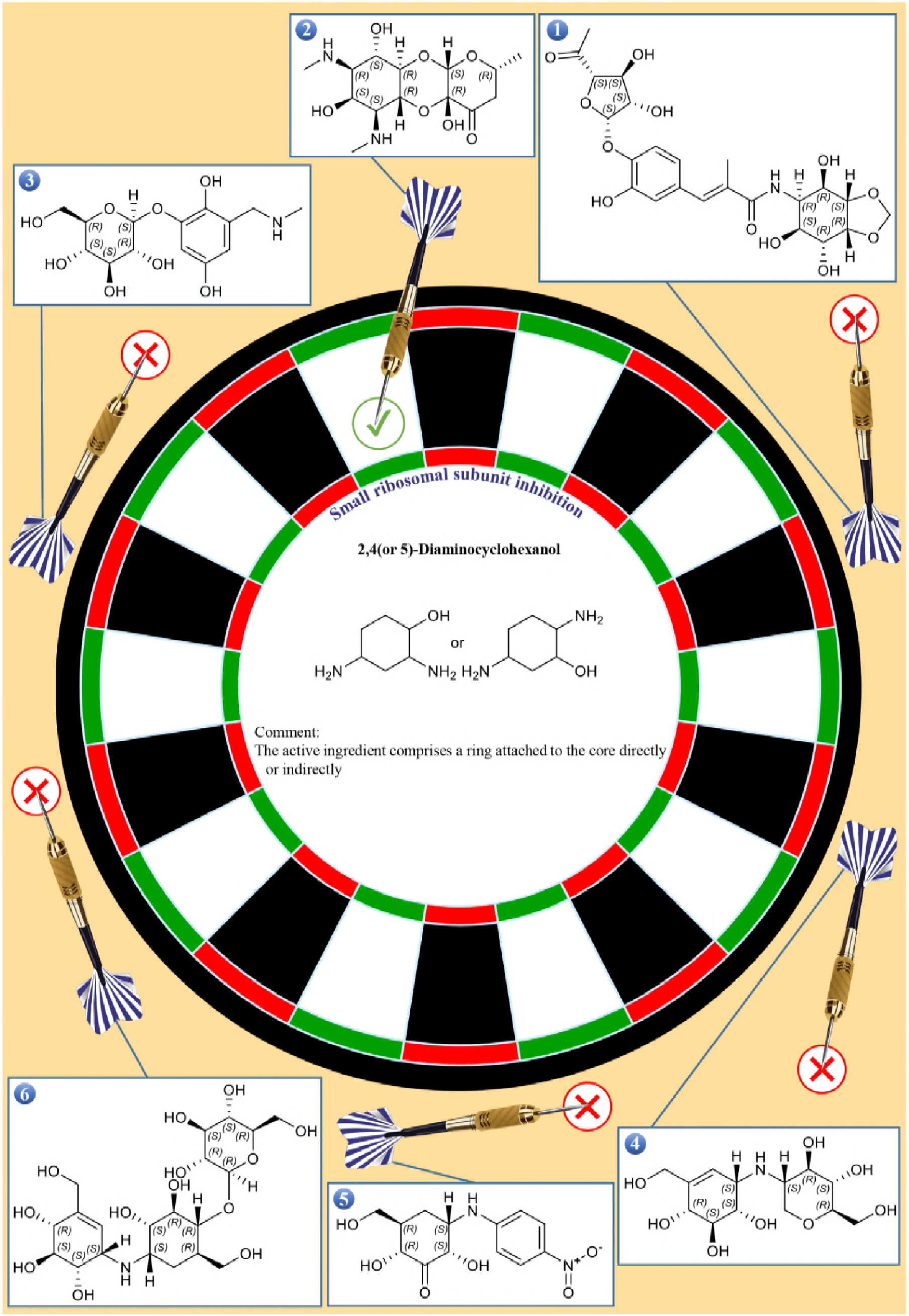
An example of identifying a bioactive molecule with the primary target of small ribosomal subunit inhibition by the minimum structure in 2,4(or 5)-diaminocyclohexanol group. Molecules 1, 3, 4, 5 and 6 don’t contain 2,4(or 5)-diaminocyclohexanol (the core). Therefore, the primary target of molecules 1, 3, 4, 5 and 6 is not small ribosomal subunit inhibition.

**Supplementary Figure 3.**
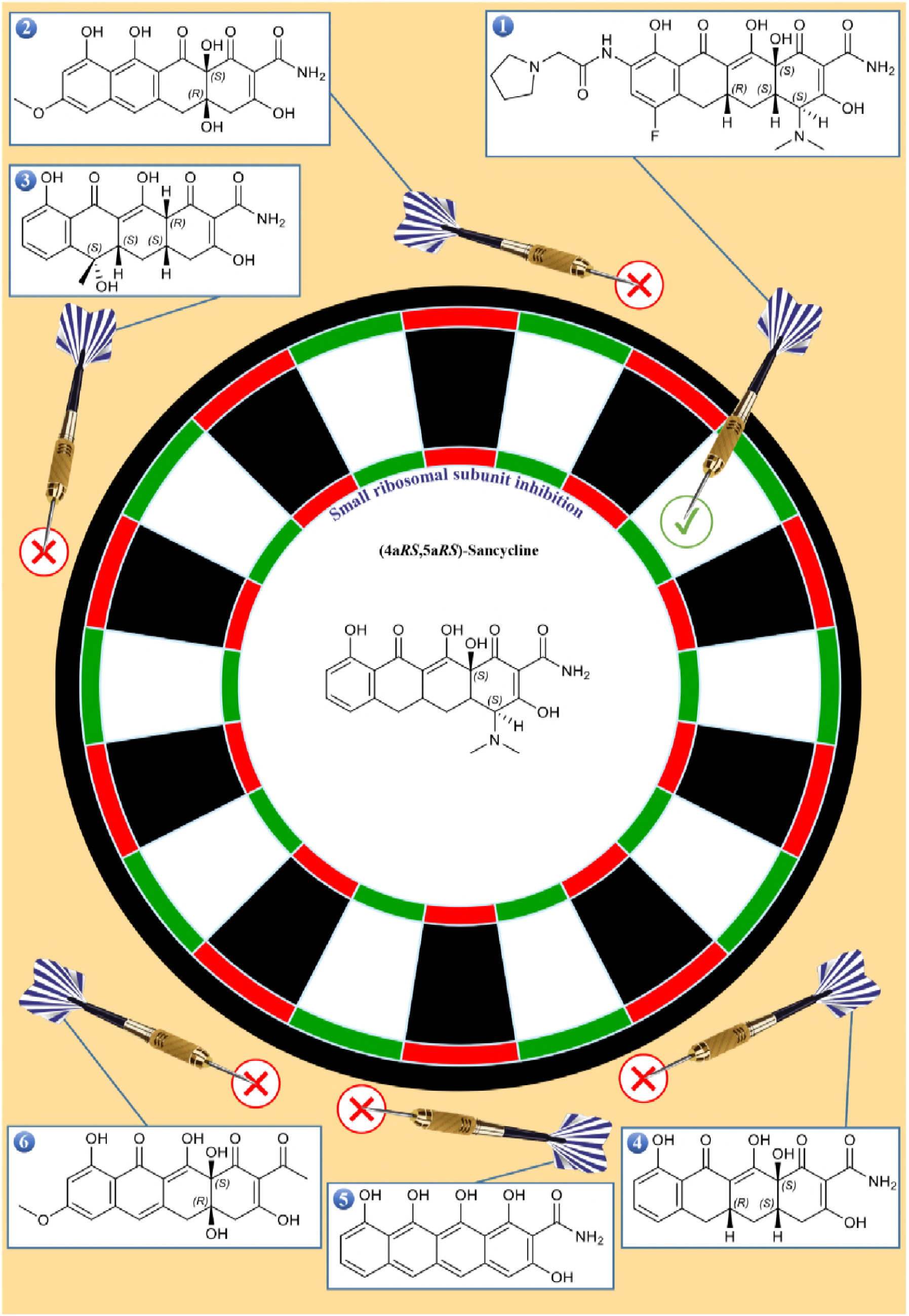
An example of identifying a bioactive molecule with the primary target of small ribosomal subunit inhibition by the minimum structure in (4a*RS*,5a*RS*)-Sancycline group. Molecules 2, 3, 4, 5 and 6 don’t contain (4a*RS*,5a*RS*)-Sancycline (the core). Therefore, the primary target of molecules 2, 3, 4, 5 and 6 is not small ribosomal subunit inhibition.

**Supplementary Figure 4.**
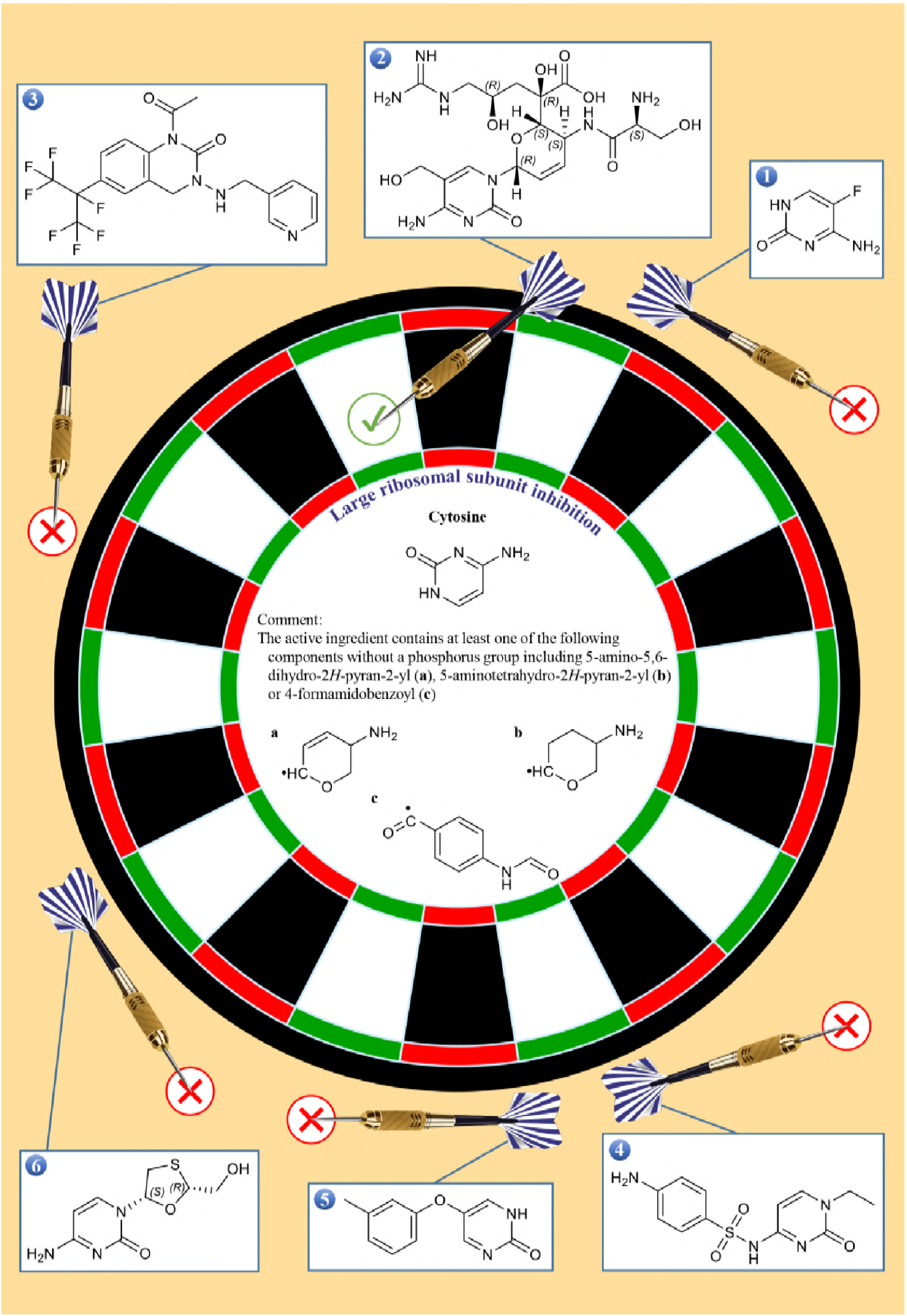
An example of identifying a bioactive molecule with the primary target of large ribosomal subunit inhibition by the minimum structure in cytosine group. Molecules 3 and 5 don’t contain cytosine (the core) and molecules 1, 4 and 6 don’t contain at least one of the following components including 5-amino-5,6-dihydro-2*H*-pyran-2-yl, 5-aminotetrahydro-2*H*-pyran-2-yl or 4-formamidobenzoyl. Therefore, the primary target of molecules 1, 3, 4, 5 and 6 is not large ribosomal subunit inhibition.

**Supplementary Figure 5.**
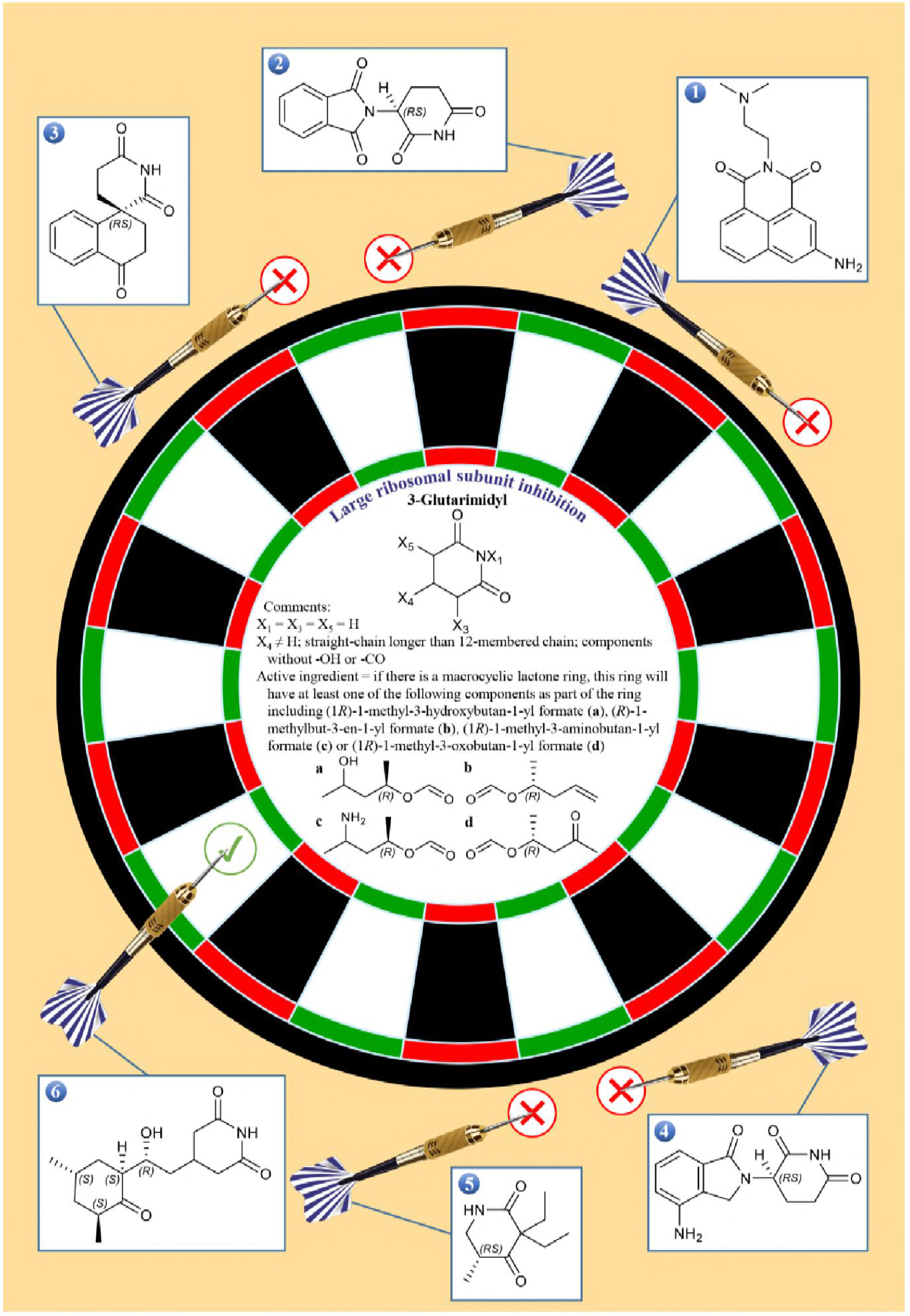
An example of identifying a bioactive molecule with the primary target of large ribosomal subunit inhibition by the minimum structure in 3-glutarimidyl group. Molecules 2, 3, 4 and 5 don’t contain 3-glutarimidyl (the core) and molecule 1 contains components other than hydrogen at X_1_, X_3_ and X_5_. Furthermore, molecule 1 contains components without -OH or -CO at X_4_. Therefore, the primary target of molecules 1, 2, 3, 4 and 5 is not large ribosomal subunit inhibition.

**Supplementary Figure 6.**
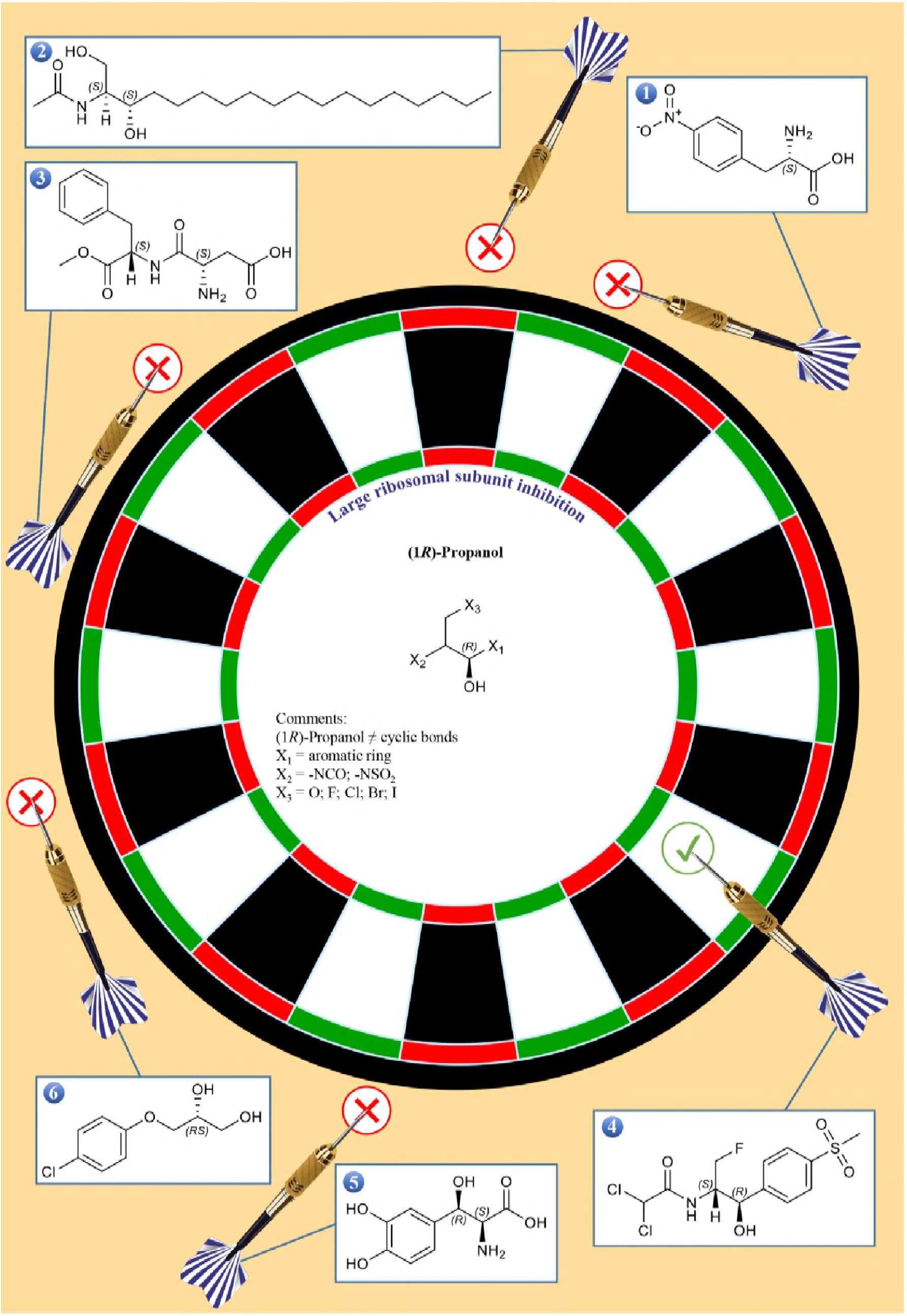
An example of identifying a bioactive molecule with the primary target of large ribosomal subunit inhibition by the minimum structure in (1*R*)-propanol group. Molecules 1, 2, 3 and 6 don’t contain (1*R*)-propanol (the core) and molecule 5 doesn’t contain -NCO or -NSO_2_ at X_2_. Therefore, the primary target of molecules 1, 2, 3, 5 and 6 is not large ribosomal subunit inhibition.

**Supplementary Figure 7.**
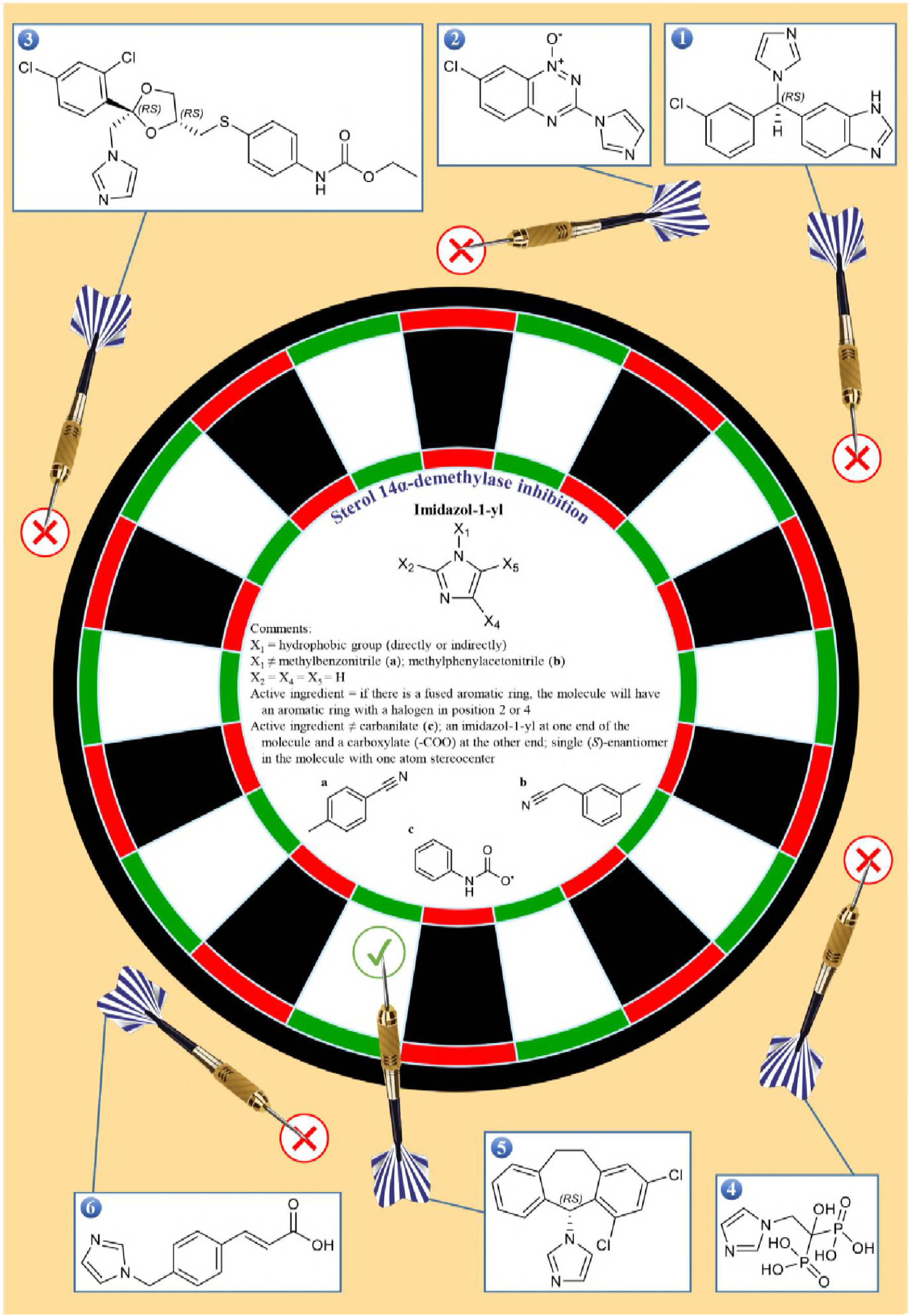
An example of identifying a bioactive molecule with the primary target of sterol 14α-demethylase inhibition by the minimum structure in imidazol-1-yl group. Molecules 1 and 2 contain a fused aromatic ring but these molecules don’t have an aromatic ring with a halogen in position 2 or 4, molecule 3 has a carbanilate, molecule 4 doesn’t contain a hydrophobic group (directly or indirectly) at X_1_ and molecule 6 contains an imidazol-1-yl at one end of the molecule and a carboxylate (-COO) at the other end. Therefore, the primary target of molecules 1, 2, 3, 4 and 6 is not sterol 14α-demethylase inhibition.

**Supplementary Figure 8.**
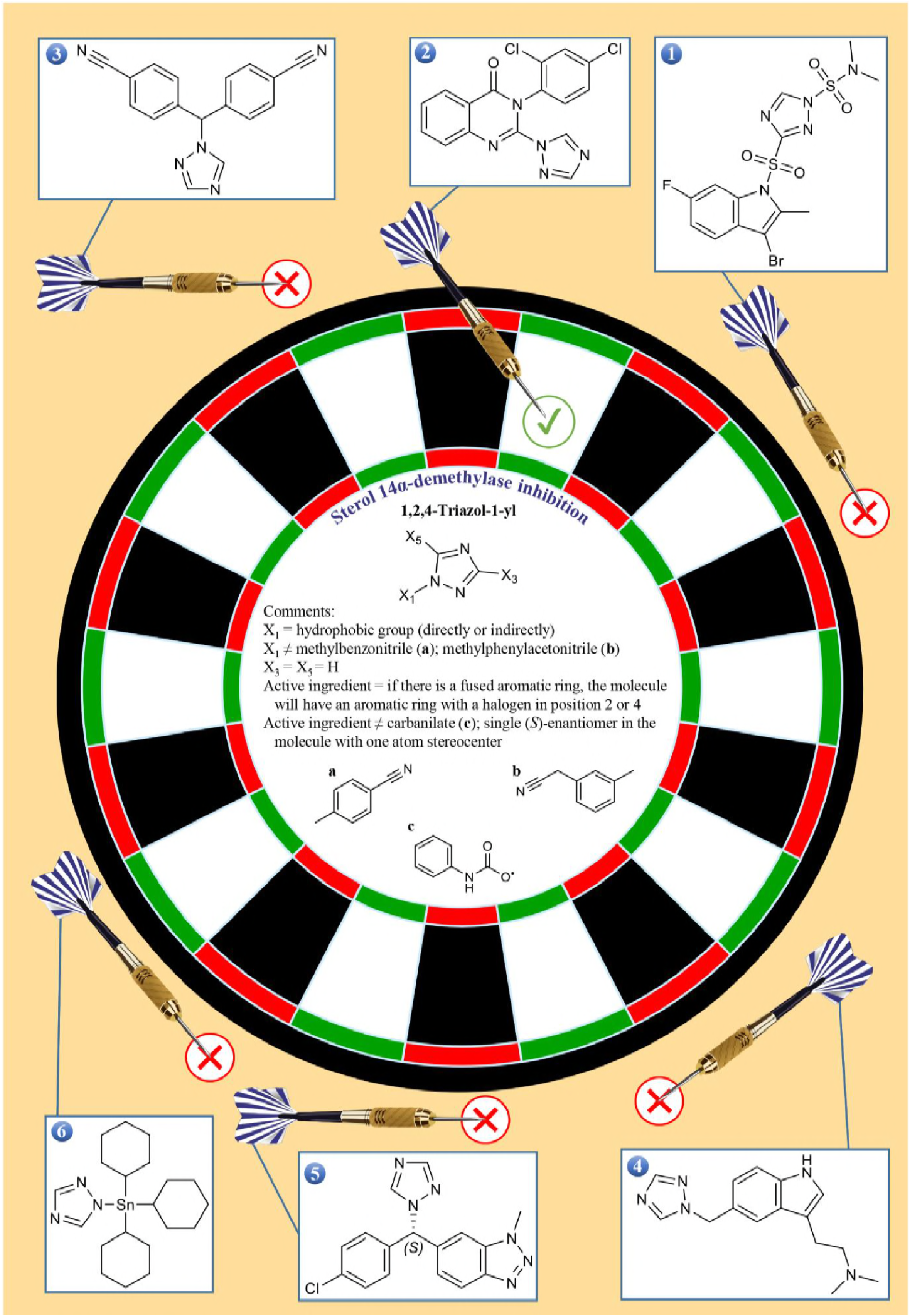
An example of identifying a bioactive molecule with the primary target of sterol 14α-demethylase inhibition by the minimum structure in 1,2,4-triazol-1-yl group. Molecule 1 contains components other than hydrogen at X_3_, molecule 3 has a methylbenzonitrile at X_1_, molecule 4 contains a fused aromatic ring but this molecule doesn’t have an aromatic ring with a halogen in position 2 or 4, molecule 5 contains one atom stereocenter and single (*S*)-enantiomer and molecules 1 and 6 don’t contain a hydrophobic group (directly or indirectly) at X_1_. Therefore, the primary target of molecules 1, 3, 4, 5 and 6 is not sterol 14α-demethylase inhibition.

**Supplementary Figure 9.**
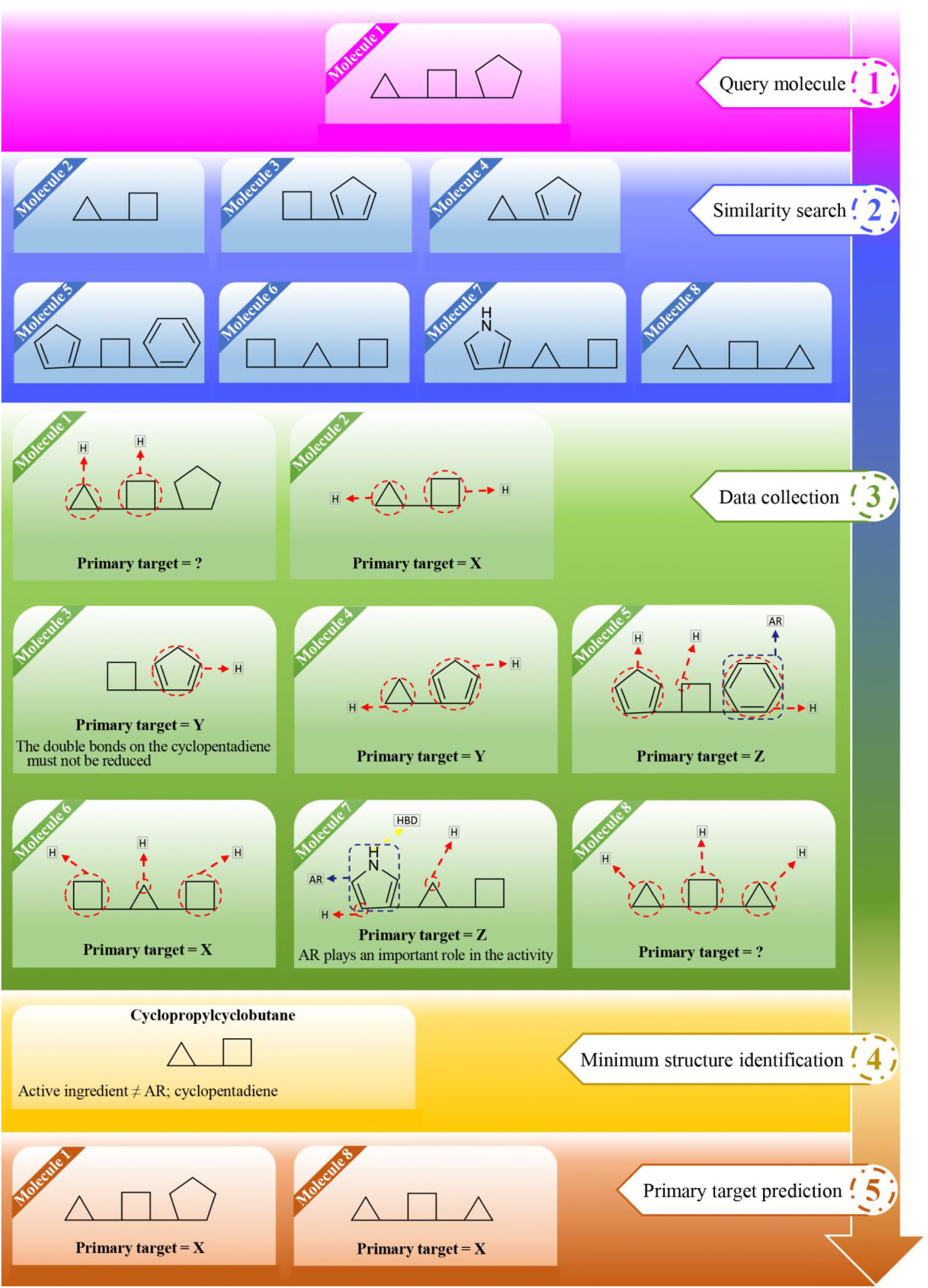
The process for predicting the primary target from chemical structure based on the minimum structure in a hypothetical example. Molecule 1 is used as the query molecule. Seven structurally related molecules to molecule 1 are found by similarity searching. Since the primary target of the query molecule (molecule 1) is unknown, information on the primary target, structure-activity relationship and pharmacophore are collected for the eight structurally related molecules. Then we consider the following assumptions: Molecules 3 and 4 have the primary target of Y. Molecule 3 consists of a square attached to a pentagon with two double bonds, and molecule 4 consists of a triangle attached to a pentagon with two double bonds. Based on information on the structure-activity relationship of molecule 3, the double bonds on the pentagon with two double bonds (cyclopentadiene) must not be reduced. Therefore, the core of these two molecules is the common and important part, named the pentagon with two double bonds (cyclopentadiene). The core of molecules 3 and 4 in the presence of square and triangle is attributed to the pentagon with two double bonds. As a result, the pentagon with two double bonds has priority over the square and the triangle in expressing the primary target of the molecule. Molecules 5 and 7 have the primary target of Z. Molecule 5 consists of a square attached to a six-membered aromatic ring (benzene) and a pentagon with two double bonds (cyclopentadiene), and molecule 7 consists of a triangle attached to a five-membered aromatic ring (pyrrole) and a square. Based on information on the structure-activity relationship of molecule 7, the aromatic ring plays an important role in the activity. Therefore, the core of these two molecules is the common and important part, named the aromatic ring (the aromatic ring in molecules 5 and 7 is characterized by LigandScout with AR). There is the pentagon with two double bonds in molecules 3, 4 and 5, but the core of molecule 5 is attributed to the aromatic ring. As a result, the aromatic ring has priority over the pentagon with two double bonds (cyclopentadiene) in expressing the primary target of the molecule. Molecules 2 and 6 have the primary target of X. Molecule 2 consists of a triangle attached to a square, and molecule 6 consists of a triangle attached to two squares. Because there is no molecule with the primary target of X having the square or the triangle alone, so the core of these two molecules is the common part, named the triangle attached to the square. The triangle attached to the square is also present in molecule 7, but the core of this molecule is attributed to the aromatic ring. As a result, the aromatic ring has priority over the triangle attached to the square in expressing the primary target of the molecule. Molecules 1 and 8 have an unknown target. Molecule 1 consists of a square attached to a triangle and a pentagon, and molecule 8 consists of a square attached to two triangles. There is also a triangle attached to a square (or a square attached to a triangle) in molecules 2, 6 and 7. In addition, the results showed that the aromatic ring and possibly the pentagon with two double bonds (cyclopentadiene) have priority over the triangle attached to the square in expressing the primary target of the molecule, but molecules 1 and 8 don’t contain any the aromatic ring or the pentagon with two double bonds. Therefore, the core of molecules 1 and 8 consists of the triangle attached to the square (cyclopropylcyclobutane) and the peripheral part does not consist the aromatic ring and the pentagon with two double bonds (cyclopentadiene). As a result, the primary target of molecules 1 and 8 is identical to the primary target of their neighbor molecules, named molecules 2 and 6, and the primary target of the two molecules is predicted X.

## Abbreviations

AR: aromatic ring
H: hydrophobic interaction
HBD: hydrogen bond donor

